# Sequential replacement of PSD95 subunits in postsynaptic supercomplexes is slowest in the cortex

**DOI:** 10.1101/2023.07.13.548866

**Authors:** Katie Morris, Edita Bulovaite, Takeshi Kaizuka, Sebastian Schnorrenberg, Candace Adams, Noboru H. Komiyama, Lorena Mendive-Tapia, Seth G. N. Grant, Mathew H. Horrocks

**Affiliations:** EaStCHEM School of Chemistry, University of Edinburgh, Edinburgh, EH9 3FJ, United Kingdom; Genes to Cognition Program, Centre for Clinical Brain Sciences, University of Edinburgh, Edinburgh, EH16 4SB, United Kingdom; EMBL Imaging Centre, European Molecular Biology Laboratory, Heidelberg, Germany; IRR Chemistry Hub, Institute for Regeneration and Repair, University of Edinburgh, Edinburgh, EH16 4UU, United Kingdom; Simons Initiative for the Developing Brain (SIDB), Centre for Discovery Brain Sciences, University of Edinburgh, Edinburgh, EH8 9XD, United Kingdom; The Patrick Wild Centre for Research into Autism, Fragile X Syndrome & Intellectual Disabilities, Centre for Discovery Brain Sciences, University of Edinburgh, Edinburgh, EH8 9XD, United Kingdom; Centre for Inflammation Research, University of Edinburgh, Edinburgh, EH16 4UU, United Kingdom

## Abstract

The concept that dimeric protein complexes in synapses can sequentially replace their subunits has been a cornerstone of Francis Crick’s 1984 hypothesis, explaining how long-term memories could be maintained in the face of short protein lifetimes. However, it is unknown whether the subunits of protein complexes that mediate memory are sequentially replaced in the brain and if this process is linked to protein lifetime. We address these issues by focusing on supercomplexes assembled by the abundant postsynaptic scaffolding protein PSD95, which plays a crucial role in memory. We used single-molecule detection, super-resolution microscopy and MINFLUX to probe the molecular composition of PSD95 supercomplexes in mice carrying genetically encoded HaloTags, eGFP and mEos2. We found a population of PSD95-containing supercomplexes comprised of two copies of PSD95, with a dominant 12.7 nm separation. Time-stamping of PSD95 subunits *in vivo* revealed that each PSD95 subunit was sequentially replaced over days and weeks. Comparison of brain regions showed subunit replacement was slowest in the cortex, where PSD95 protein lifetime is longest. Our findings reveal that protein supercomplexes within the postsynaptic density can be maintained by gradual replacement of individual subunits providing a mechanism for stable maintenance of their organization. Moreover, we extend Crick’s model by suggesting that synapses with slow subunit replacement of protein supercomplexes and long protein lifetimes are specialized for long-term memory storage and that these synapses are highly enriched in superficial layers of the cortex where long-term memories are stored.

---

Synapses in the central nervous system allow the transmission of information between neurons and are the site at which memories are formed and stored. The vast majority of synapses in the mammalian brain employ glutamate as the neurotransmitter, which is released from the presynaptic terminal on axons and diffuses onto the postsynaptic terminal on dendrites where it binds glutamate receptors (Kandel, E.R. et al., 2021). Activating glutamate receptors triggers the biochemical changes in the postsynaptic terminal that ultimately encode memories. The proteome of the postsynaptic termini of excitatory synapses is highly complicated and comprises over 1000 proteins representing many structural and functional classes of molecules (Sorokina et al., 2021). Biochemical studies show that all these proteins are organized into supramolecular assemblies of complexes and supercomplexes (complexes of complexes) (Collins et al., 2006; Collins and Grant, 2007; Fernández et al., 2017, 2009; Frank and Grant, 2017; Frank et al., 2016; Husi et al., 2000; Husi and Grant, 2001; Nithianantharajah et al., 2013; Pocklington et al., 2006). The best described complexes are the ionotropic glutamate receptors known as NMDA (N-methyl-D-aspartate) and AMPA (α-amino-3-hydroxy-5-methyl-4-isoxazolepropionic acid) receptors, which are formed from four membrane-spanning subunits that together produce a ligand-gated ion channel (Greger et al., 2017; Hansen et al., 2018). Major supercomplexes are those formed by the scaffolding protein PSD95 (Cho et al., 1992), which can assemble various complexes including NMDA and AMPA receptors, other ion channels, adhesion proteins and signaling proteins (Fernández et al., 2009; Frank and Grant, 2017; Frank et al., 2016; Husi et al., 2000). Mice and humans carrying mutations in PSD95 show profound learning and memory deficits (Migaud et al., 1998; Nithianantharajah et al., 2013), indicating that the formation and function of supercomplexes is crucial for storage of information in the brain.

Although it is widely accepted that rapid biochemical changes in the postsynaptic termini of excitatory synapses underlie the initial encoding of memories, the mechanisms that control the duration of memories and rate of forgetting remain poorly understood. There has been a long-standing quest to identify molecular changes that could persist for the duration of the memory. This became a central issue in memory research in 1984 as a result of an influential article by Francis Crick (Crick, 1984). He reasoned that the biochemical changes in synapses would be erased by protein turnover, and that because some memories must have a longer lifetime than synaptic proteins, he postulated that there must be molecular mechanisms that perpetuated the initial biochemical changes. Toward this, he posited that dimeric complexes of proteins may be modified during learning and gradually replace their subunits in a way that would allow the ‘molecular memory’ in a preexisting subunit to be transferred to the new subunit in a mixed complex of old and new subunits. As a result of this, much attention has focused on the enzymatic complex known as calcium/calmodulin-dependent protein kinase II (CamKII), which is comprised of multiple subunits that have the capacity for autophosphorylation and persistent activation (Bayer and Schulman, 2019; Bhattacharyya et al., 2016; Hell, 2014; Stratton et al., 2013). Experiments using recombinant proteins *in vitro* show that holoenzymes of CamKII can exchange subunits, which in principle could be a mechanism for perpetuation of molecular memories (Bhattacharyya et al., 2020, 2016; Singh and Bhalla, 2018; Stratton et al., 2013) . However, it is unknown if old subunits are sequentially replaced by new subunits *in vivo* in CamkII complexes, or indeed any other complexes in synapses.

PSD95 affords an opportunity to revisit Crick’s hypotheses for several reasons. First, biochemical studies of PSD95 supercomplexes isolated from the mouse brain reveal there is a family of supercomplexes where members all appear to comprise two copies of PSD95 and different combinations of interacting proteins (Frank and Grant, 2017; Frank et al., 2016). Second, excitatory synapses show high diversity arising from the differential distribution of protein complexes (Cizeron et al., 2020; Zhu et al., 2018) and differential lifetimes of PSD95 (Bulovaite et al., 2022). Those synapses with longest PSD95 lifetimes were referred to as Long Protein Lifetime (LPL) synapses, and a brainwide analysis of their distribution showed they were most highly enriched in the cortex where long-term memories are stored. These findings raise the following questions: are the two copies of PSD95 in supercomplexes sequentially replaced, or does the dimeric complex degrade and rebuild from two new copies of PSD95? Secondly, if there is a sequential replacement of PSD95 subunits in supercomplexes, is the rate of this replacement related to the protein lifetime?

To address these questions, we have used single-molecule imaging of PSD95-containing supercomplexes isolated directly from the mouse brain. Using lines of mice that contain genetically-encoded tags fused to the carboxyl terminus of endogenous PSD95 (Broadhead et al., 2016; Bulovaite et al., 2022; Zhu et al., 2018), we prepared protein extracts from the brain and imaged supercomplexes using Total Internal Reflection Fluorescence (TIRF) microscopy. We counted the number of PSD95 copies in each supercomplex, and found that the majority contain two units. We next measured the mean distance between the labels using MINFLUX, which has a spatial resolution of 1-5 nm, and showed that the average distance between the fluorophores is 12.7 nm. Using mice that express HaloTag fused to PSD95 (Bulovaite et al., 2022), we injected a fluorescent ligand into the tail vein, which efficiently labels PSD95 in brain synapses, and by extracting supercomplexes at time points after injection, we show that individual PSD95 subunits are sequentially replaced, in line with Crick’s hypothesis. Finally, we ask if the rate of subunit exchange is influenced by PSD95 protein lifetime by comparing brain regions with different PSD95 lifetimes. We found that the rate of exchange is slowest in the cortex where the protein lifetime is longest. Our results, which are the first visualization of individual synaptic supercomplexes and their constituent proteins, show that synaptic scaffold proteins that play a crucial role in memory are organized as dimers in supercomplexes, and are maintained by sequential replacement of individual subunits over extended periods of time, and that this process is linked to the rate of protein turnover in the regions of the brain involved with long-term memory storage.

## Imaging individual PSD95 supercomplexes isolated from mouse brain

Single-molecule and super-resolution (SR) approaches enable the heterogeneity in molecular complexes and supercomplexes to be distinguished (Jain et al., 2011; Saleeb et al., 2023; Szymborska et al., 2013), and therefore provide an invaluable tool for studying synaptic molecular supercomplexes isolated from brain homogenate. Forebrains from PSD95-GFP homozygous mice were dissected and homogenized as described (Fernández et al., 2009) (**Supplementary Methods and Materials**) (**Figure 1a**). The supercomplexes were subsequently diluted and immobilized on a glass coverslip, and imaged on a TIRF microscope (**Figure 1bi**) (**Supplementary Methods and Materials**). Although this technique is diffraction-limited, the number of fluorescent proteins present in each supercomplex can be quantified by the stepwise photobleaching of each fluorophore (Dalton et al., 2016; Leake et al., 2006). The number of photobleaching steps per diffraction-limited spot was determined for a population of >6,000 supercomplexes across three biological repeats (**Figure 1bii**, with examples of one-step and multi-step photobleaching traces presented in **Figure 1ciii**). On average, there were 1.6 PSD95 proteins per PSD95-containing supercomplex; however, taking advantage of our ability to characterize individual supercomplexes, we found that 63% contained one PSD95 protein, 24% two PSD95 proteins, and 13% more than two PSD95 proteins. Given that not all eGFP will fold correctly (typical *in vitro* refolding yields for GFP are 50–60% (Battistutta et al., 2000; Reid and Flynn, 1997; Ward and Bokman, 1982)), this suggests that there is an abundant population of PSD95 supercomplexes that contain two PSD95 proteins. We validated our approach using purified eGFP, demonstrating that the vast majority of eGFP was monomeric (**Figure S1**).

**Figure 1.**
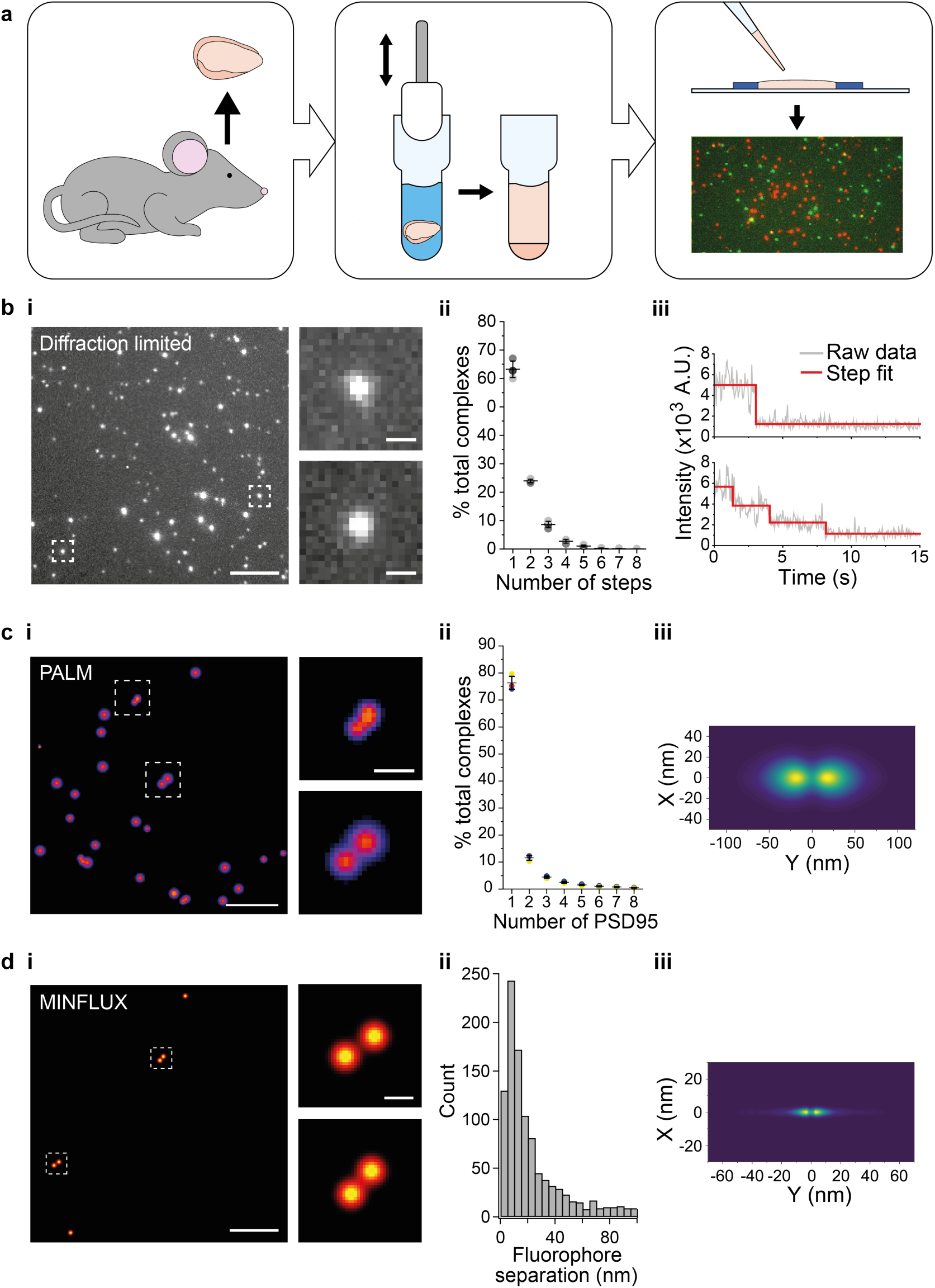
Stoichiometry and spatial arrangement of PSD95 within individual supercomplexes. a) Brain containing either endogenously-tagged PSD95-GFP or PSD95-mEoS was extracted from the genetically-modified mouse, and the forebrain was homogenized to solubilize PSD95-containing supercomplexes. PSD95 supercomplexes were immobilized on glass coverslips and imaged using single-molecule and super-resolution approaches. bi) Individual PSD95-GFP supercomplexes (boxed) were imaged using TIRF microscopy. Scale bar = 5 µm, inset scale bar = 500 nm. Photobleaching step counting revealed a distribution of PSD95 stoichiometries (bii) (10,178 PSD95-GFP photobleaching steps were counted across 6,461 supercomplexes). Representative intensity traces with fits shown in biii. ci) Example PALM images of individual PSD95 supercomplexes. Scale bar = 500 nm, inset scale bar = 50 nm. cii) Subsequent analysis revealed that PSD95 exists at a range of stoichiometries within the supercomplexes (132,929 PSD95-mEoS molecules were detected in 82,501 individual supercomplexes). Plots show mean ± SD, n = 3 biological repeats. ciii) Class averaging of the dimer population shows a distinct separation between the PSD95 proteins within the supercomplexes (class average of 9,743 supercomplexes). di) Example MINFLUX images of PSD95 supercomplexes. Scale bar = 100 nm, inset scale bar = 10 nm. dii) Analysis of the supercomplexes containing two PSD95 molecules (1011 supercomplexes) showed a distribution of PSD95 separation distances. diii) Class averaging of this population revealed two peaks separated by 12.7 nm.

To further verify the PSD95 stoichiometry in supercomplexes, we generated brain homogenate from PSD95-mEos heterozygous mice, and following immobilization, we imaged the supercomplexes using photoactivated localization microscopy (PALM) (Betzig et al., 2006). Each PSD95-mEos molecule within the supercomplexes was localized with a mean precision of 21.8 ± 6.5 nm (mean ± SD, n = 132,929 localizations across 3 biological repeats). Fourier ring correlation (Nieuwenhuizen et al., 2013) revealed that the acquired images had a mean resolution of 30 ± 4 nm (mean ± SD, n = 3 biological repeats). This enabled the spatial relationship between individual PSD95 molecules to be deduced. Examples of super-resolved supercomplexes are shown in **Figure 1ci**. The number of PSD95 molecules per supercomplex was quantified using custom-written code (**Supplementary Methods and Materials**), and the distribution of stoichiometries is shown in **Figure 1cii**. Over 130,000 PSD95-mEoS proteins were localized in 82,501 supercomplexes across three biological repeats. 76 ± 2% of the supercomplexes contained only one PSD95-mEoS protein, 12 ± 1% contained two PSD95-mEoS proteins, and the remainder (12 ± 2%) of the clusters contained more than two PSD95-mEoS proteins. Given that only half of the PSD95 proteins are fused to mEoS2 in heterozygous mice, and that not all mEoS2 will fold correctly, these results confirm the findings from the photobleaching analysis, suggesting that supercomplexes contain PSD95 protein at a range of stoichiometries, with the majority containing two or fewer. We also identified multimers in wild-type PSD95 supercomplexes using a mix of orthogonally labeled PSD95 nanobodies (**Figure S2**).

We next determined the mean distance between PSD95 molecules by class averaging the 9,743 supercomplexes that contained two PSD95-mEoS proteins, revealing one dominant separation with a mean distance of 37.8 nm between PSD95 molecules (**Figure 1ciii**). As this separation distance is close to the spatial resolution that we were able to achieve using PALM, we further analyzed the supercomplexes using MINFLUX, which can attain a spatial resolution of 1-5 nm. We immobilized supercomplexes from brain homogenate containing PSD95-GFP and added a GFP-nanobody tagged with Alexa Fluor 647 (**Supplementary Methods and Materials.** Details of MINFLUX imaging sequences are provided in **Table S1**). Example MINFLUX images are shown in **Figure 1di**. We were able to detect PSD95 within the supercomplexes at a range of stoichiometries, and selected for measurement those that contained two units of PSD95 (1011 supercomplexes). Class averaging the distribution of individual distances measured between two PSD95 molecules in the supercomplexes (**Figure 1dii**) shows two clear peaks separated by 12.7 nm (**Figure 1diii**).

## Sequential replacement of PSD95 within supercomplexes

We have previously measured the rate of PSD95 turnover across the mouse brain at single-synapse resolution, demonstrating that excitatory synapses have a wide range of protein lifetimes extending from a few hours to several months (Bulovaite et al., 2022). These findings led us to ask whether protein turnover could occur within individual intact molecular supercomplexes, or if the whole supercomplex needs to be replaced with new protein.

Injecting the cell- and blood-brain barrier-permeable fluorescent ligand Silicon-Rhodamine-Halo (SiR-Halo) into the tail vein of 3-month-old PSD95-HaloTag homozygous mice labels all of the PSD95-HaloTag (**Figure 2a**) (Bulovaite et al., 2022). Because SiR-Halo forms a covalent bond with the PSD95-HaloTag, the persistence of labeling over time after injection reports whether PSD95 has been replaced in individual supercomplexes. Brain tissue was obtained at 6 hours (day-0) or 7 days (day-7) post SiR-Halo injection. The homogenized forebrain tissue from each mouse was then incubated with a second HaloTag ligand, Alexa Fluor 488-Halo (AF488-Halo), to label any new PSD95-HaloTag protein generated after the earlier SiR-Halo injection. This allowed us to quantify the levels of PSD95 protein turned over in 7 days. Supercomplexes containing only SiR-Halo represent those in which no PSD95 replacement had occurred, whereas those with only AF488-Halo are either new supercomplexes or those in which all PSD95 has been replaced. A coincident signal of SiR-Halo and AF488-Halo represents supercomplexes in which a proportion of the PSD95 has been replaced (**Figure 2a**).

**Figure 2.**
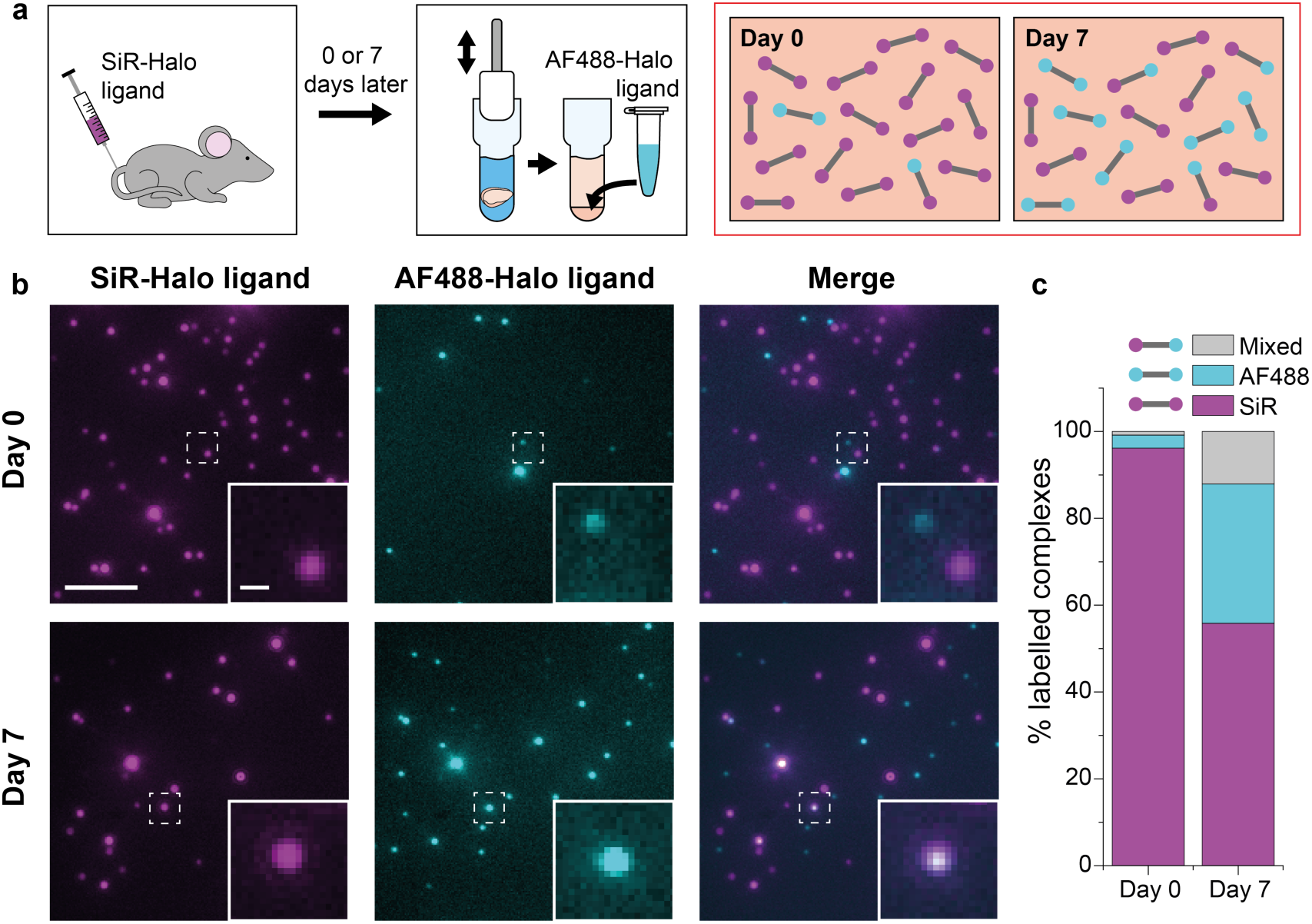
PSD95 turnover within supercomplexes in mouse brain homogenate. a) PSD95-HaloTag homozygous mice were injected with SiR-Halo ligand and culled 6 hours (day-0) or 7 days (day-7) post injection. The forebrains were homogenized and post-hoc stained with AF488-Halo ligand to saturate remaining binding sites. At day-0, the vast majority of PSD95 is labeled with SiR-Halo only. After 7 days of protein turnover, three populations of supercomplex are possible: SiR-Halo only, AF488-Halo only, both fluorophores. b) Images of supercomplexes from homogenate at day-0 and day-7, showing increased AF488-Halo:SiR-Halo ratios at day-7, with increased coincidence. Scale bar is 5 μm and 500 nm in the zoom. c) Quantified percentages of supercomplexes labeled only with SiR-Halo, AF488-Halo, or both. At day-0, 96% of supercomplexes were labeled with SiR-Halo only, indicating saturation of PSD95-HaloTag binding sites by injection. At day-7, this had decreased to 56%, with expansion of the AF488-Halo and co-labeled populations, indicating that PSD95 protein turnover had occurred over the 7 days.

Example diffraction-limited images of the labeled PSD95 supercomplexes at the two time points are shown in **Figure 2b**. Most of the supercomplexes observed at day-0 contain only SiR-Halo, with few having AF488-Halo, and negligible coincidence. At day-7, fewer SiR-Halo supercomplexes were observed, indicating that some of the PSD95 protein present at the time of injection was degraded. An increase in the number of AF488-Halo-labeled supercomplexes indicates that the degraded protein has been replaced with new, label-free protein. An increase in coincidence can be seen in the merge of the two images. These results are shown quantitatively in **Figure 2c** (see also **Figure S3**). Approximately 40% of the old PSD95 protein is replaced by new protein between day-0 and day-7. Of the 13,710 supercomplexes analyzed at day-0, 96% of the supercomplexes were labeled with only SiR-Halo, indicating that the HaloTag binding sites were saturated by injection. At day-7, 56% of the 15,391 supercomplexes analyzed were labeled with only SiR-Halo. 32% of the supercomplexes were labeled with only AF488-Halo, indicating all copies of PSD95 within these supercomplexes had been turned over in the 7-day period. Interestingly, 12% of the supercomplexes were labeled with both AF488-Halo and SiR-Halo, indicating that some supercomplexes (hereafter referred to as ‘mixed supercomplexes’) contain both old and new protein. This indicates that supercomplexes can exchange old copies of PSD95 for new.

**Figure 3.**
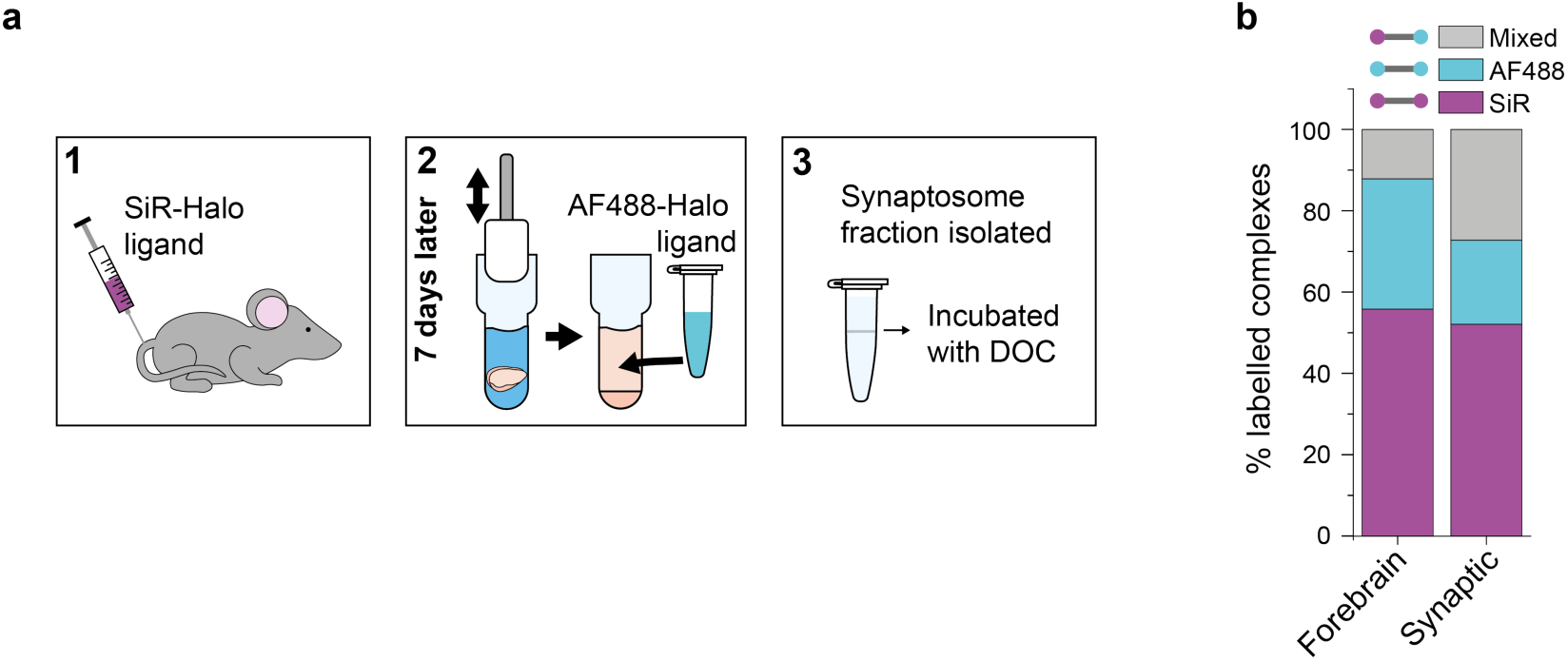
Protein turnover in synaptic and total forebrain PSD95 supercomplexes. a) SiR-Halo ligand was injected into the tail vein of three 3-month-old PSD95-HaloTag knock-in mice. 7 days later, the forebrain was extracted and synaptosomes isolated. AF488-HaloTag ligand was incorporated during the homogenization process. DOC, deoxycholate. b) Comparison of populations of PSD95 supercomplexes in the total forebrain and synaptic fraction.

**Figure 4.**
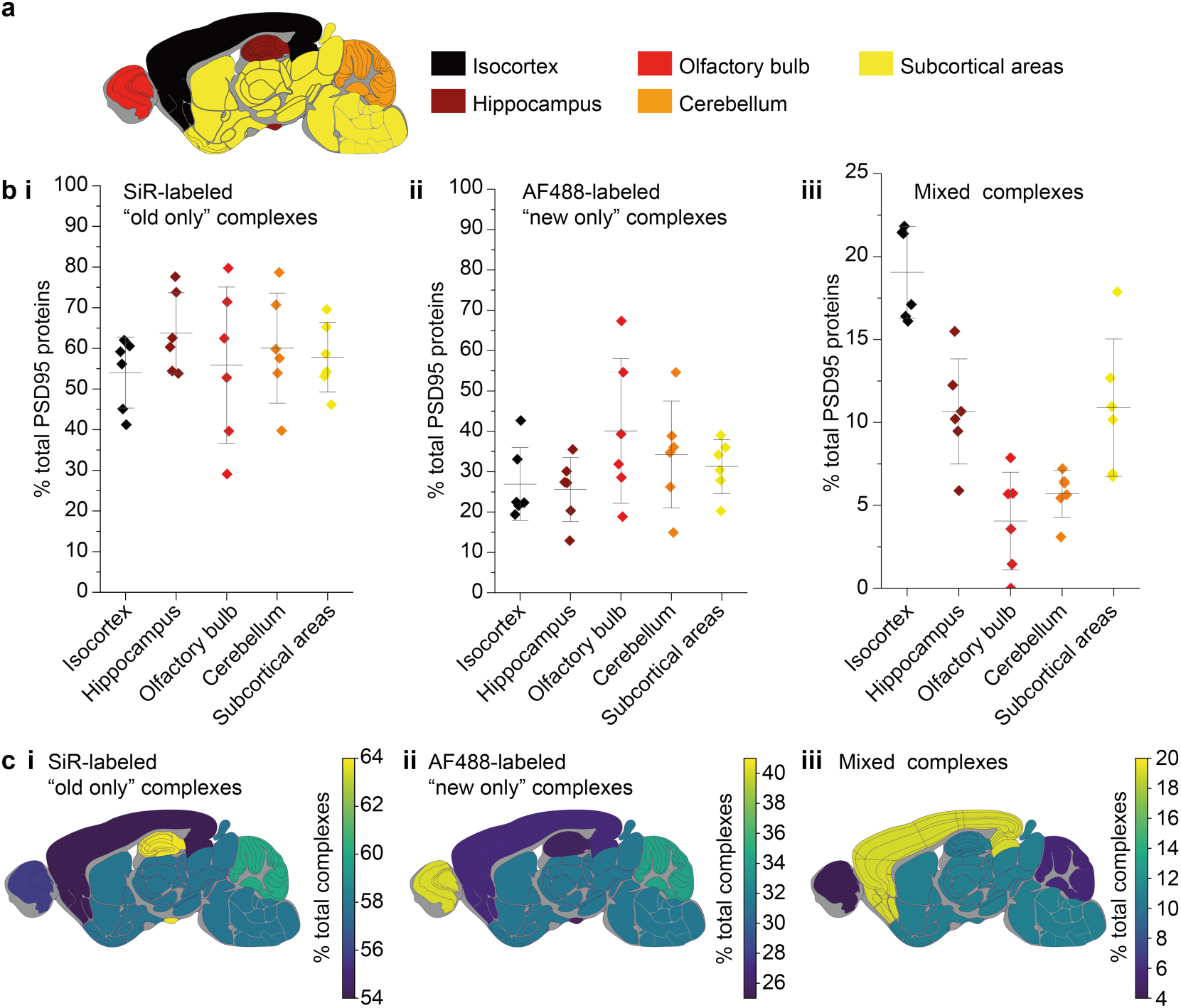
Differences in PSD95 turnover between mouse brain regions. a) Mouse brains were dissected into five broad regions. b) The percentage of total PSD95 imaged contained in SiR-labeled (i), AF488-labeled (ii) and mixed (iii) supercomplexes. Error bars show the SD of 6 biological repeats. c) The percentage of SiR-labeled (i), AF488-labeled (ii) and mixed (iii) PSD95 supercomplexes in each region of the brain. For statistical significance between the mixed complexes, see **Supplementary Methods and Materials** (Table S2).

## Comparison of synaptic PSD95 with total PSD95

PSD95 is synthesized in the neuronal cytosol and transported into synapses where it is concentrated in the postsynaptic density. We asked whether the exchange of PSD95 in supercomplexes differs between the synapse and the cytosol by comparing the synaptic (synaptosome) and total forebrains of mice injected with SiR-Halo at day-7 (**Supplementary Methods and Materials**). Supercomplexes were extracted from either the synaptosome fraction or the whole forebrain of SiR-Halo-injected mice and incubated with AF488-Halo and imaged using TIRF microscopy (**Figure 3a**).

Whereas the level of SiR-Halo-only supercomplexes is similar for both the total forebrain and synaptic fractions (56% and 52%, respectively), the percentage of AF488-Halo-only supercomplexes is lower in the synaptic fraction (21%, versus 32% in the total forebrain homogenate) and there is a greater fraction of mixed supercomplexes (27%, versus 12% in the total forebrain homogenate) (**Figure 3b**). This suggests that while the overall rate of PSD95 turnover is similar in synapses compared with the overall forebrain, there is a greater fraction of supercomplexes that retain at least one old PSD95 protein, underlying their importance in maintaining the overall molecular state of the postsynaptic density.

## Slowest exchange of PSD95 in supercomplexes from the cortex

Because the lifetime of PSD95 is longer in cortical regions than other brain regions (Bulovaite et al., 2022), we hypothesized that this region may have different rates of exchange of PSD95 within supercomplexes. The brains from six 3-month-old PSD95-HaloTag mice, culled 7 days after SiR-HaloTag ligand injection, were dissected into five major regions (isocortex, hippocampus, olfactory bulb, cerebellum and subcortex), and homogenized to extract the supercomplexes. After incubation with AF488-HaloTag ligand, to label new PSD95-Halo proteins, they were imaged using TIRF microscopy (**Figure 4a**).

In total, 60,798 supercomplexes were analyzed in the isocortex, 15,842 in the hippocampus, 3,148 in the olfactory bulb, 36,126 in the cerebellum and 28,339 in the subcortical areas. Strikingly, the isocortex contained the highest percentage of mixed supercomplexes (19 ± 3%, mean ± SD), and the olfactory bulb the lowest (4 ± 3%, mean ± SD) (**Figure 4b,c**). Correspondingly, the region with the highest percentage of new complexes was the olfactory bulb (40 ± 16%, mean ± SD), whereas the isocortex had one of the lowest percentages (27 ± 8%, mean ± SD) (**Figure 4b,c**) (statistical significances shown in **Table S2**). This pattern was also seen in 3-week-old mice (**Supplementary Methods and Materials**, **Figure S4**).

The correlation between the percentage of PSD95 proteins contained in old, new and mixed supercomplexes in each brain region was compared with the previously published half-life of PSD95 in the same regions (**Supplementary Methods and Materials**, **Figure S5**) (Bulovaite et al., 2022). Although there is limited correlation between the percentage of PSD95 proteins contained in old or new supercomplexes and the half-lives of PSD95 puncta in each region, there is a statistically significant correlation for mixed supercomplexes, with a Pearson’s correlation test value of R = 0.95 at a significance level of P = 0.01.

## Discussion

The application of single-molecule detection, SR microscopy, and MINFLUX to study genetically-encoded fluorescently-tagged PSD95 has enabled us to reveal the structure of individual postsynaptic supercomplexes at the nanometer length scale. Although we have previously used biochemical approaches to demonstrate that, on average, two copies of PSD95 exist in supercomplexes in bulk protein extracts (Frank and Grant, 2017), this study represents the first direct observation of PSD95 in these megadalton supercomplexes at single-molecule resolution. Using multiple single-molecule and SR microscopy techniques to image the brain homogenate from three mouse models (PSD95-HaloTag, PSD95-GFP, PSD95-mEos), we have robustly cross-validated our approach for studying the organization of PSD95 in individual supercomplexes. Unlike ensemble averaging techniques, a major advantage of our approach is the ability to observe and quantify hundreds of thousands of PSD95 molecules in tens of thousands of individual supercomplexes, allowing the stoichiometry in each one to be determined, as well as the distances between the PSD95 molecules. Furthermore, by tagging PSD95-HaloTag with specific Halo ligands *in vivo*, followed by imaging at different time-points, our approach allows the lifetime of PSD95 in individual supercomplexes to be measured. Dissection of brain regions prior to homogenization and imaging also enables our methods to be deployed for generating maps of structure and lifetime of supercomplexes in different brain areas.

PSD95 is one of the most abundant synaptic proteins and is localized beneath the postsynaptic membrane of excitatory synapses in a structure known as the postsynaptic density. Nanoscale imaging of PSD95 in synapses using PALM and STED reveals individual synapses differ in their PSD95 content and spatial organization. For example, in the mouse hippocampus and cortex there are populations of synapses with single or multiple 40 nm ‘nanoclusters’ and larger contiguous structures (Broadhead et al., 2016; Masch et al., 2018). Receptors, including AMPARs have also been shown to distribute into such nanoclusters (MacGillavry et al., 2013) and these have been shown to align to clusters at the presynapse via ‘nanocolumns’, demonstrating their functional role (Tang et al., 2016). Bulk protein extracts reveal that all PSD95 is assembled into 1-3 MDa supercomplexes (Frank and Grant, 2017; Frank et al., 2016; Husi et al., 2000) indicating that they are the building blocks of the nanoscale structures observed with SR techniques. We now show that this synaptic heterogeneity extends to the individual supercomplex level, and demonstrate that while the majority of supercomplexes contain two or fewer PSD95 molecules, others contain more. With the enhanced resolution of MINFLUX, we were able to measure the distances between individual PSD95 molecules within the supercomplexes and find that, rather than being fixed, there is considerable variation; however, class averaging showed a dominant separation distance of 12.7 nm. This could reflect a difference in the molecular make-up of each supercomplex. Indeed, supercomplexes are composed of dozens of proteins (Collins et al., 2006; Fernández et al., 2009; Frank and Grant, 2017; Husi et al., 2000) and biochemical characterization of their composition from different brain regions show different supercomplexes populate synapses in different regions (Frank and Grant, 2017). Moreover, brainwide synaptome mapping of the proteins in individual synapses show synapse diversity arises from their constituent complexes and supercomplexes (Cizeron et al., 2020; Tomas-Roca et al., 2022; Zhu et al., 2018). Two recent studies have provided insights into the spacing between PSD95 molecules. MINFLUX revealed a mean nearest neighbor distance of approximately 7 nm (Gürth et al., 2023), while one-step nanoscale expansion microscopy found a preferred spacing of 8-9 nm (Shaib et al., 2023). These measurements, taken within the PSD, may represent distances between neighboring supercomplexes rather than within a single supercomplex, as reported here. Furthermore, in our study, the slightly larger measurements may be due to the size of the GFP molecules (∼3 nm) and the nanobodies used, which could contribute to an increased observed separation between PSD95 molecules

The maintenance of synaptic structure at the molecular level is necessary to maintain physiological stability of brain circuits. Brainwide mapping of the spatial distribution of PSD95-expressing synapses across the lifespan shows that between 3-9 months of age the remarkable synapse diversity and its organization into the synaptome architecture is very stable (Cizeron et al., 2020). However, when we measured the lifetime of PSD95 during this age window, we found that the vast majority of PSD95 is replaced every few weeks (Bulovaite et al., 2022). Thus, the synaptome architecture, which is comprised of molecularly diverse synapses, is stable despite its constituent proteins being replaced. Our present findings offer an explanation for how this stability can be maintained: the protein supercomplexes which are the building blocks of the synaptome architecture are not removed and replaced in toto, but are maintained by the sequential replacement of constituents including core scaffolding proteins.

## Memory maintenance by sequential subunit replacement and protein lifetime

In the synaptome theory of behavior (Grant, 2018; Zhu et al., 2018), representations, memories and behavioral programs are encoded in the synaptome architecture, and thus its stability is required to maintain these functional outputs. The stability conferred on the synaptome architecture by the sequential replacement of subunits in supercomplexes offers a stability mechanism. However, it is also necessary to have plasticity to learn new things and to forget unnecessary information. We propose that synapse diversity resolves this dilemma: there are some synapses that are very stable, and others that are less stable and more plastic. Our present findings indicate that the most stable synapses are those with long-protein lifetime in the cortex, and that these synapses undergo slow replacement of their supercomplex subunits. The processes of subunit replacement and protein turnover are synergistic mechanisms that produce stability of supramolecular entities, which together with synapse diversity can result in synapses with a range of memory durations and plasticity potential.

This model fits with Crick’s basic framework in that it involves dimeric proteins that sequentially exchange their subunits. Our findings extend his model in that we show a link between the protein lifetime and the rate of subunit exchange. At the time Crick made his postulate, the rate of protein turnover in the brain and synapses was unknown, and until recently, there was no evidence that synapses might differ in their protein lifetime. The surprising finding that some synapses can maintain copies of PSD95 for months draws into question Crick’s assumption that protein lifetimes are much shorter than memories, because most memories are forgotten within a short period (days to weeks) in mice.

It is interesting to speculate how the exchange of PSD95 subunits could lead to the perpetuation of a molecular and behavioral memory. There are families of PSD95 supercomplexes that contain different proteins and particular interacting proteins may be well-suited to modifying the newly arrived subunit and altering its properties. The concentration of PSD95 supercomplexes is highest in the postsynaptic terminal and this environment may facilitate exchange of subunits between supercomplexes, consistent with our results. Some synapses such as the long-protein lifetime synapses that are enriched in the cortex may be better suited to the exchange process and transmission of modifications. Indeed, it may be that the environment of these synapses facilitates other complexes and supercomplexes (such as CamKII holoenzymes) to exchange and/or replace subunits and that our observations with PSD95 reflect a more widespread and general phenomenon. Our findings and approaches provide a platform for addressing these issues as well as providing a new model of memory duration.

## AUTHOR INFORMATION

Corresponding Authors

Seth G. N. Grant - Genes to Cognition Program, Centre for Clinical Brain Sciences, University of Edinburgh, Edinburgh EH16 4SB, United Kingdom.

Mathew H. Horrocks - EaStCHEM School of Chemistry, The University of Edinburgh, Edinburgh EH9 3FJ, United Kingdom.

## Author Contributions

MHH and SGNG conceptualized the idea and designed the experiments. KM, EB, TK, CA, SS, NHK, and LMT performed experiments and analyzed the data. The manuscript was written by KM, SGNG and MHH with contribution from all authors. All authors have given approval to the final version of the manuscript.

## Acknowledgments

1. J. Dorrens, G. Varga, E. Robson for management of mouse colony and genotyping. C. Davey for editing.

## ABBREVIATIONS

SR: super-resolution
PALM: photoactivation localization microscopy
MINFLUX: MINimal photon FLUXes
TIRF: total internal reflection fluorescence
GFP: green fluorescent protein
LPL: long protein lifetime
SPL: short protein lifetime

