## Supplementary Information for "Sequential replacement of PSD95 subunits in postsynaptic supercomplexes is slowest in the cortex"

| | |
|---|---|
| Materials and Methods | |
| Table S1 | MINFLUX sequences used. |
| Figure S1 | Stoichiometry of eGFP determined using photobleaching step count analysis. |
| Figure S2 | Multimers of PSD95 in wild-type supercomplexes. |
| Figure S3 | Absolute number graphs for Figure 2 c, d and e. |
| Figure S4 | Characterization of PSD95 turnover in 3-week-old mice. |
| Figure S5 | Comparisons between the density based PSD95 puncta half-life as published in Bulovaite et al. |
| Statistical analysis for Figure 4 | |
| Table S2 | Post-hoc t-tests to locate the source of significance between the means of the five brain regions analyzed (mixed complexes). |
| References | |

MATERIALS AND METHODS

### Preparation of mouse brain homogenates

Homogenates were prepared from dissected brains or whole mouse forebrains, as described previously (Frank et al., 2016). Deoxycholate (DOC) extraction buffer (1% sodium deoxycholate, 50 mM tris(hydroxymethyl)aminomethane (tris) pH 9.0, 50 mM sodium fluoride, 20 μM zinc chloride, 1 mM sodium ortho-vanadate, 2 mM 4-(2-aminoethyl)-benzene-sulfonyl fluoride and 1 Complete Protease Inhibitor Cocktail tablet (Roche, Germany) per 50 mL) was prepared and stored on ice prior to homogenization.

For whole brain homogenization, each forebrain was added to 5 mL DOC buffer and homogenized using 20 strokes with a 5 mL capacity Teflon homogenizer. The homogenate was stored on ice for 1 hour, homogenizing again using 20 strokes at 30 minutes. The homogenate was centrifuged at 50,000 g for 30 minutes at 4$^{o}$C. The supernatant was separated from the pellet and contained the PSD95 supercomplexes.

For dissected brain homogenization, each dissected brain region was added to 0.6 mL DOC buffer. Homogenization and centrifugation were performed as above for whole brain specimens, except that pellet pestles (Fisher Scientific) were used instead of a teflon homogenizer to homogenize the tissue.

### Preparation of synaptosome fractions

Adult (5-month-old) PSD95-HaloTag mice were injected with HaloTag ligand conjugated with SiR as described previously (Bulovaite et al., 2022). Seven days after injection, the mice were sacrificed with cervical dislocation and the forebrain was dissected, briefly rinsed with ice-cold PBS, frozen with liquid nitrogen, and stored at –80℃ before use. For synaptosome preparation, homogenization of tissue was performed by 12 strokes with Teflon-glass homogenizer in Homogenization buffer (0.32 M sucrose, 1 mM HEPES pH 7.4, and Complete EDTA-free Protease Inhibitor Cocktail (Merck)). Brain homogenate was centrifuged at 1,400 g for 10 min at 4℃ to obtain the pellet and the supernatant fraction. The pellet fraction was resuspended in Homogenization buffer with 3 strokes of the homogenizer and centrifuged at 700 g for 10 min at 4℃. The supernatant of the first and second centrifugation was pooled as an S1 fraction and subjected to centrifugation at 14,000 g for 10 min at 4℃. The resulting pellet was resuspended with Homogenization buffer (P2 fraction) and centrifuged in a sucrose density gradient (0.85/1.0/1.2 M) for 2 hours at 82,500 g. The fraction between 1.0 M and 1.2 M was collected and used as synaptosome.

#### Dissection of mouse brain regions

Mice were culled by cervical dislocation and decapitated. The brains were removed from the skull and quickly washed with ice-cold PBS. Brains were dissected on ice, on a plastic plate covered with 3MM filter paper soaked in ice-cold PBS. The cerebellum and olfactory bulbs were removed first, then the hippocampus and cortex were isolated from the rest of the brain. Brain samples were snap frozen in liquid nitrogen and stored at -80°C.

#### Protein turnover measurements

For the protein turnover experiments, homozygous PSD95$^{HaloTag/HaloTag}$ knock-in mice were injected with 200 µL 1.5 mM SiR-Halo ligand diluted in saline and Pluronic F-127 (20% in DMSO). As a control, some injected mice were culled 6 hours post injection to provide a 0-day timepoint with maximum saturation. The remainder of the mice were culled 7 days post injection. Their brains were extracted and prepared as forebrain or region-specific homogenates as described above. Prior to imaging, the turnover homogenates were incubated with 10 µM AF488-Halo ligand in a 1:1 volumetric ratio for 1 hour at 4$^{o}$C.

**TIRF microscopy**

All diffraction-limited and PALM experiments were conducted on a home-built TIRF microscope described previously(de Moliner et al., 2023). Briefly, collimated laser light at 405 nm (Cobolt MLD 405–250 Diode Laser System, Cobalt, Sweden), 488 nm (Cobolt Fandango-300 DPSS Laser System, Cobalt, Sweden), 561 nm (Cobolt DPL561-100 DPSS Laser System, Cobalt, Sweden), and 638 nm (Cobolt MLD Series 638-140 Diode Laser System, Cobolt AB, Solna, Sweden) was aligned and directed parallel to the optical axis at the edge of a 1.49 NA TIRF objective (CFI Apochromat TIRF 60XC Oil), mounted on an inverted Nikon TI2 microscope. The microscope was fitted with a perfect focus system to auto-correct the z-stage drift during imaging. Fluorescence collected by the same objective was separated from the returning TIR beam by a dichroic mirror Di01-R405/488/561/635 (Semrock, Rochester, NY, USA), and was passed through appropriate filters (488 nm: BLP01-488R-25, FF01-525/30-25; 561 nm: LP02-568RS, FF01-587/35; 638 nm: FF01-432/515/595/730-25, LP02-647RU-25, Semrock, Rochester, NY, USA). Fluorescence was then passed through a 2.5× beam expander and recorded on an EMCCD camera (Delta Evolve 512, Photometrics) operating in frame transfer mode (EMGain = 11.5 e-/ADU and 250 ADU/photon). Each pixel was 103 nm in length. Images were recorded with an exposure time of 50 ms. The microscope was automated using the open-source microscopy platform Micromanager. Borosilicate glass coverslips (20 × 20 mm, VWR International) were cleaned using an Ar plasma cleaner (Zepto, Diener) for 30 min to remove any fluorescent residues. Frame-Seal slide chambers (9 × 9 $mm^2$, Bio-Rad) were affixed to the glass to create a well in which samples (100 µL) were added. All samples in the wells were washed three times with PBS prior to imaging.

For diffraction-limited photobleaching analysis of GFP, the neat homogenate was diluted 1:100 in PBS and irradiated and imaged with 488 nm (68 W $cm^{-2}$, 25 s), ensuring all molecules were photobleached. For diffraction-limited photobleaching analysis of SiR and AF488, whole forebrain homogenate and DOC-treated synaptosomes were diluted 1:100 in PBS. Homogenates from dissected brain regions were diluted 1:1000 to 1:10,000 in PBS. The samples were first irradiated and imaged with 638 nm light (850 W $cm^{-2}$, 25 s), followed by 488 nm light (110 W $cm^{-2}$, 25 s), ensuring all molecules were photobleached. For PALM imaging, the samples were diluted 1:100 in PBS and illuminated with 15 cycles of 405 nm (75 W $cm^{-2}$, 1 s) and 561 nm (1500 W $cm^{-2}$, 10 s) irradiation until all molecules were photobleached. In preparation for imaging, samples were incubated on the glass surface for 30 seconds prior to washing three times with 0.02 µm filtered PBS.

**Coincidence analysis**

Coincidence analysis was carried out using a custom written MATLAB script. Initially, all spots in the diffraction limited images were detected using the Find Maxima function in ImageJ. The prominences used varied depending upon the fluorophores and the power of the lasers, but generally were ~500-1000. The locations of all spots in both channels were loaded into MATLAB. The distances between the $n^{th}$ spot in the first channel and all spots in the second channel were calculated. Any spots with a separation distance less than the channel offset parameter (2 pixels in this work) were classed as being ‘coincident’. Finally, the number of spots in the first channel, the number of spots in the second channel, and the number of coincident spots were output by the script.

All scripts used in this analysis are available at: DOI: 10.5281/zenodo.8059239

**Photobleaching analysis**

Photobleaching data were collected using the TIRF microscope described above, and analyzed using a published approach (Chappard et al., 2023; Choi et al., 2022). The intensity of the emitted fluorescence from all molecules was tracked over a period of 25 seconds. The intensity traces from each molecule were extracted by finding spots in the first frame of the image stack using the ImageJ Find Maxima function. The intensity at each spot location was then measured in all frames of the stack using a custom written ImageJ macro. Chung-Kennedy filtering was performed on the resulting intensity traces using a custom written MATLAB script, with a shuttling window size of 12. The Chung-Kennedy filter (Chung and Kennedy, 1991) shuttles two windows along the data set either side of each data point. The output of the Chung-Kennedy filter is a weighted average of the mean of the two windows. The weighting shifts the output value towards the mean of the window with the lowest variance such that noise is reduced, but discontinuities in the data do not become blurred during the filtering process.

To detect photobleaching steps in the filtered traces, an approximate differential of the filtered data was calculated. Peaks in the differentiated intensity traces indicated a sharp change in gradient in the intensity trace and thus the position of the photobleaching steps. A peak threshold of 75 was applied to the differentiated intensity traces to extract the location of the steps. The validity of each of the steps was determined by calculating the ratio of individual step height to the local regional variance. This calculation returned a t-statistic for each potential step. A t-threshold of 0.1 was applied to distinguish true steps from false steps. True steps have large ratios of step height to local regional variance and thus larger t-statistics. False steps have smaller ratios of step height to local regional variance and thus smaller t-statistics. Once the location of each step was known and the validity confirmed, a step function was generated as a fit to the raw step data.

All scripts used in this analysis are available at: DOI: 10.5281/zenodo.7118670

**PALM analysis**

The data were preliminarily analyzed using the Peak Fit function of the GDSC SMLM ImageJ plugin to output super-resolved localizations of the blinking fluorophores. A signal strength threshold of 20 was used, along with a precision threshold of 40 nm. Following this, the drift was corrected for using the Drift Calculator function in the GDSC SMLM ImageJ plugin. Once the localizations were extracted, a custom written MATLAB clustering script was used. The clustering script sorted all localizations by precision from low precision to high precision. To correct for multiple blinks emanating from the same fluorophore, the script consolidated all localizations within the precision of another localization into one object. The distances between all objects were then calculated. Objects separated by less than 160 nm were grouped into clusters. The number of objects in each cluster were counted. Clusters containing one object were defined as ‘monomeric’, clusters containing two objects were defined as ‘dimeric’, etc. Information pertaining to the clusters (number of objects, x-y position of objects, average precision) was output as a text file.

Class averaging was performed on the class of clusters containing two objects by another custom written MATLAB script. The script aligns all dimeric clusters parallel to the x-axis and centers them about the midpoint between the two objects. The objects are plotted as width = precision, and the resulting density at each x-y position is calculated to give the surface plot showing the class average.

All scripts used in this analysis are available at: DOI: 10.5281/zenodo.7993694

**MINFLUX**

MINFLUX imaging was conducted on an Abberior 3D-MINFLUX microscope (Abberior Instruments, Göttingen, Germany) equipped with a 100x oil immersion objective lens (UPL SAPO100XO/1.4, Olympus, Tokyo, Japan). MINFLUX imaging of Alexa 647 was performed using a 642 nm CW excitation laser at 22.6 μ/cm$^2$ in the first MINLUX iteration. Laser powers were measured at the position of the objective lens back focal plane using a Thorlabs PM100D power meter equipped with a S120C sensor head. Fluorescence signal from Alexa 647 was detected using two avalanche photodiodes (SPCM-AQRH-13, Excelitas Technologies, Mississauga, Canada) with a detection range of 650 – 685 nm for the first detector and 685 – 760 nm for the second detector channel (detected photons were summed). The pinhole was set to a size corresponding to 0.78 airy units for all imaging experiments. The microscope was operated by Abberior Imspector software (version 16.3.13924-m2112). The build-in stabilization system was used to minimize drift of the sample for the duration of the measurement. Therefore, scattering from 200 nm gold nanoparticles (Nanopartz, Cat# A11-200-CIT-DIH-1-10, USA) which were deposited on the coverslip surface was used as a positional reference for the active sample stabilization.

For MINFLUX imaging of PSD95-GFP, borosilicate glass coverslips (No. 1.5H, round, 24 mm diameter, product # 117640, Marienfeld, Germany) were Ar plasma cleaned for 15 min to remove any fluorescent residues. Next, coverslips were incubated with PSD95 protein lysate (1:100 in PBS) at room temperature for 1 hour. After three washing steps with PBS, coverslips were incubated with 200 nm goldparticles (Nanopartz, Cat# A11-200-CIT-DIH-1-10, USA) diluted 1:10 in PBS for 10 minutes. To remove goldparticles which did not attach, coverslips were washed three times with PBS and subsequently blocked using 0.5% BSA in PBS for 20 minutes at room temperature. Next, PSD95GFP was stained using an anti-GFP nanobody coupled with Alexa 647 (FluoTag®-X4 anti-GFP-A647, Nanotag, Germany) diluted 1:100 in 0.5% BSA in PBS and incubated overnight at 4°C. Next day, the sample was washed three times with PBS and mounted in GLOX buffer (50 mM Tris, 10 mM NaCl, 10% glucose (w/v), 500 µg/ml glucose oxidase, 40 µg/ml catalase, pH 8.0) supplemented with 20 mM MEA on cavity slides and sealed using Twinsil (Picodent, Germany). For 2D-MINFLUX imaging of PSD95-GFP, the MINFLUX sequence with 5 iterations was used (Table S1).

**Table S1. MINFLUX Iterations.**

| | 1st | 2nd | 3rd | 4th | 5th |
|---|---|---|---|---|---|
| **L size [nm]** | 300 | 300 | 150 | 75 | 40 |
| **TCP - pattern** | Hexagon | Hexagon | Hexagon | Hexagon | Hexagon |
| **Minimum number of collected photons** | 100 | 150 | 100 | 100 | 150 |
| **Laser power factor** | 1x | 1x | 2x | 4x | 6x |
| **TCP dwell time [ms]** | 1 | 1 | 1 | 1 | 1 |
| **CFR** | x | 0.5 | x | 0.8 | x |
| **Background threshold [kHz]** | 15 | 15 | 15 | 15 | 15 |

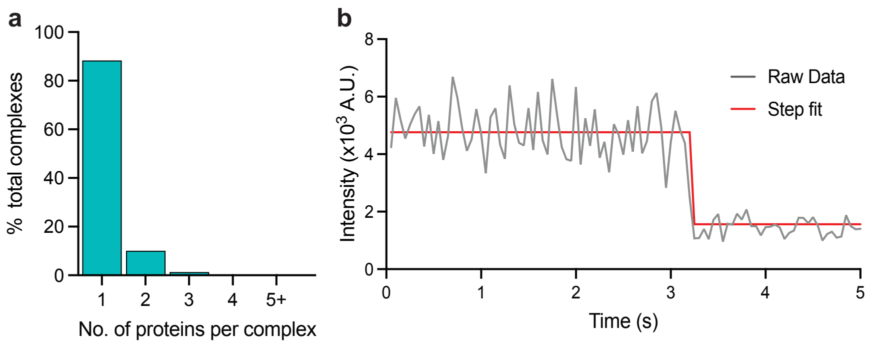


**Figure S1. Stoichiometry of eGFP determined using photobleaching step count analysis. a)** Stoichiometries of eGFP determined using the same approach as that for eGFP-tagged PSD95. **b)** Example intensity trace with fitting.

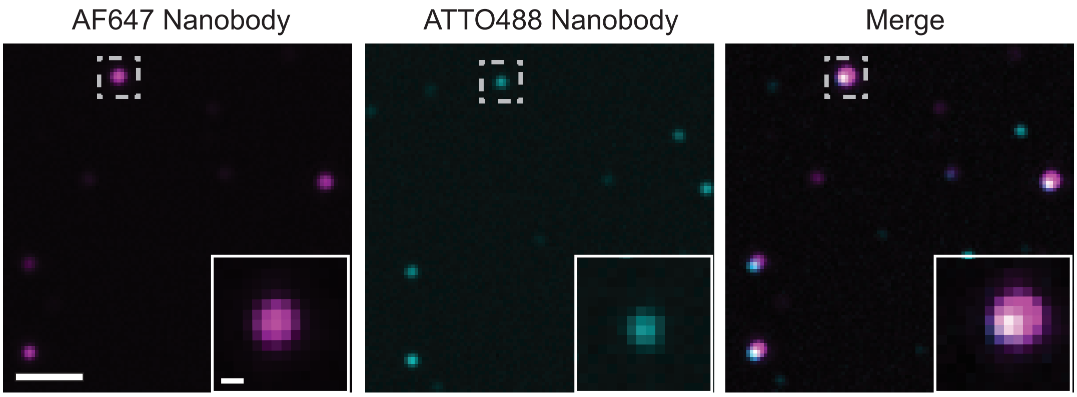


**Figure S2. Multimers of PSD95 in wild-type supercomplexes**. Homogenate extracted from wild-type mice was processed using the same procedure as that for PSD95-eGFP mice. It was subsequently diluted 1:1000 and 100 µl of this solution was incubated on a plasma-cleaned coverslip for 30 minutes. The coverslip was washed three times with PBS and a mix of 2.5 pM AF647-tagged and 2.5 pM ATTO488-tagged FluoTag®-X2 anti-PSD95 nanobody (NanoTag Biotechnologies) was added to the coverslip (total volume 100 µl) and incubated for 30 minutes. Following washing three times with PBS, the sample was imaged on the TIRF microscope with 638 nm and 488 nm excitation. Coincident spots represent supercomplexes containing multiple PSD95 units. Scale bar = 2 µm, inset scale bar = 250 nm.

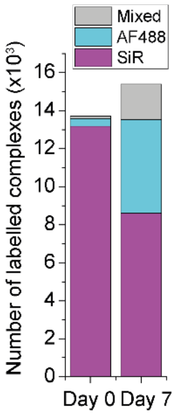


**Figure S3. Absolute number graphs for Figure 2c.** Number of labeled supercomplexes at day-0 and day-7 containing only SiR-labeled 'old' protein (SIR),

AF488-labeled ‘new’ protein (AF488), or both ‘mixed’. In total, 13,710 supercomplexes were analyzed at day-0 and 15,391 supercomplexes were analyzed at day-7.

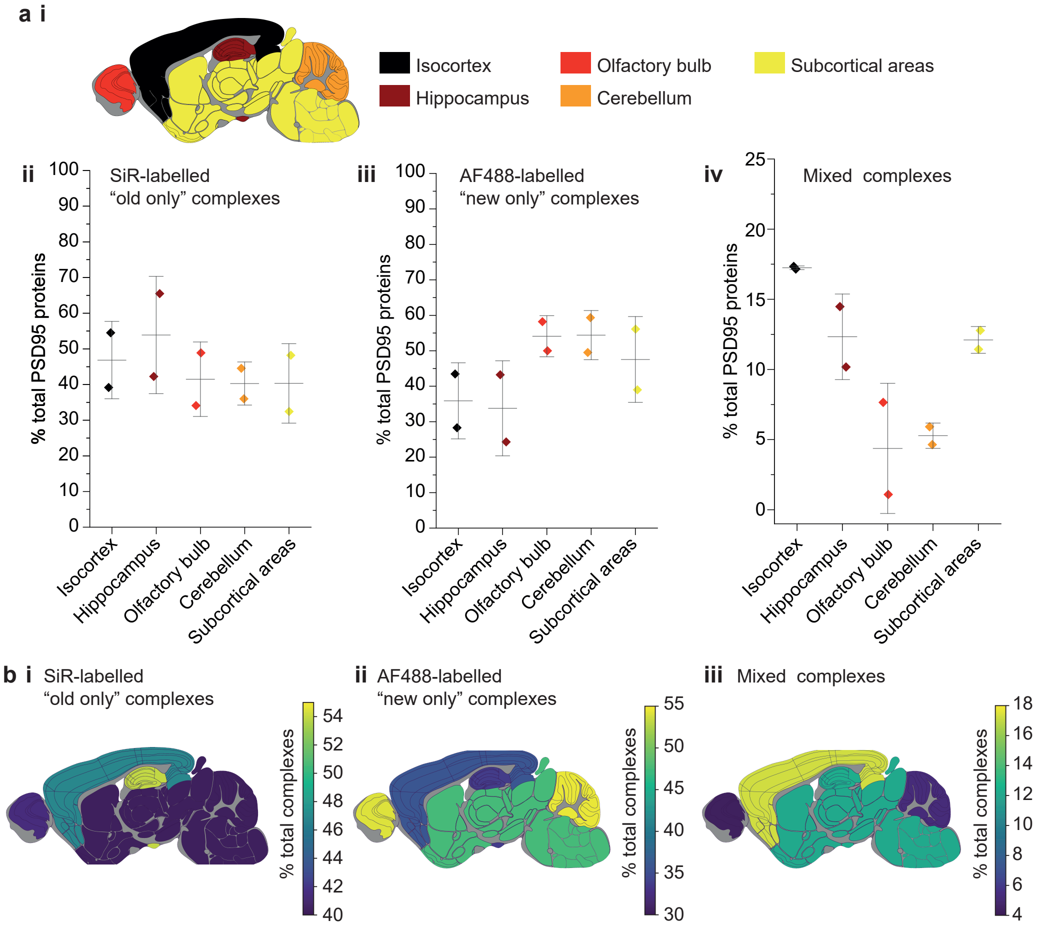


**Figure S4. Characterization of PSD95 turnover in 3-week-old mice. a)** Key showing the five dissected brain regions (**i**). Percentage of total PSD95 proteins in old-only (**ii**), new-only (**iii**), and mixed (**iv**) supercomplexes (mean ± SD, n = 3). **b)** Mouse brain region heatmaps showing the same data as in a ii-iv.

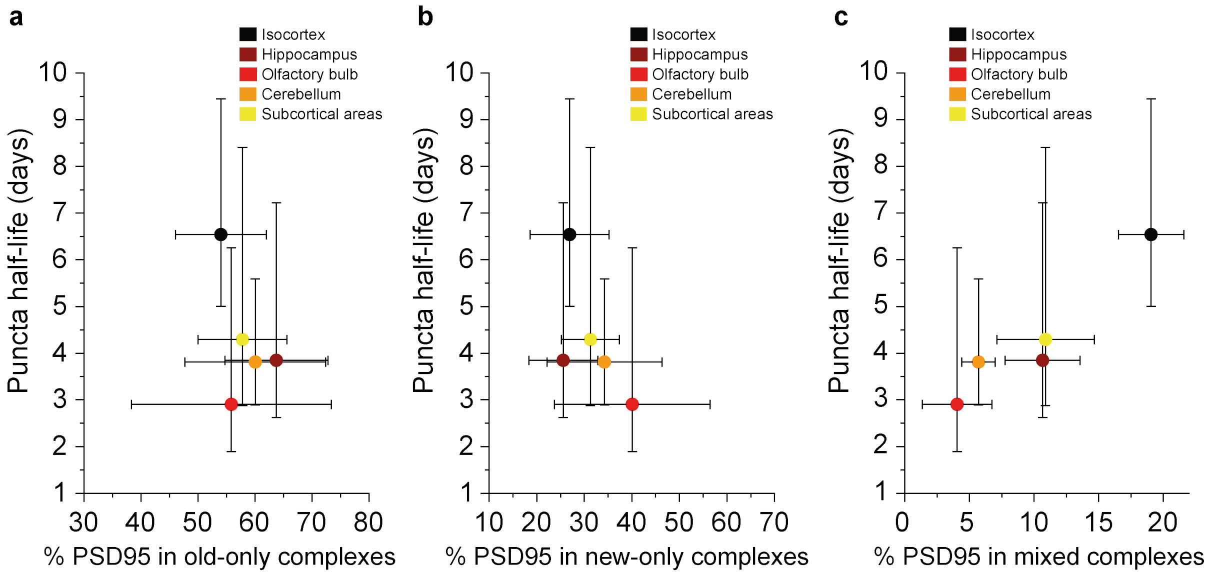


**Figure S5. Comparison between the density based PSD95 puncta half-life and old, new and mixed proteins in supercomplexes.** The percentage of complexes that contain **a**) old-only, **b**) new-only or **c**) mixed old and new PSD95 (mean $\pm$ SD for % PSD95, and $\pm$ 95% CI for half-life, n = 3). There is no statistically significant correlation in a or b. However, R = 0.9536 and P = 0.01 from a Pearson's correlation test, indicating a significant positive correlation between the percentage of PSD95 in mixed supercomplexes and PSD95 puncta half-life.

### Statistical analysis for Figure 4

ANOVA tests for differences between the five brain region means was carried out for plots bi, bii and biii in Figure 4. The ANOVA tests returned P-values of 0.71, 0.23 and $5.5 \times 10^{-8}$ respectively. This indicates that there is no statistical significance in plots bi and bii, but there are statistical differences between some of the means in plot biii. Post-hoc t-tests were carried out to identify the source of the significance. The P-values for comparisons between all regions are shown in Table S2.

**Table S2.** Post-hoc t-tests to locate the source of significance between the means of the mixed supercomplexes in the five brain regions analyzed. Only P-values less than 0.05 are shown. There are high levels of significance between the means of the isocortex and all other regions.

| | **Isocortex** | **Hippocampus** | **Olfactory bulb** | **Cerebellum** | **Subcortex** |
|---|---|---|---|---|---|
| **Isocortex** | | $6.4 \times 10^{-4}$ | $3.8 \times 10^{-6}$ | $1.5 \times 10^{-5}$ | $3.1 \times 10^{-3}$ |
| **Hippocampus** | | | $3.8 \times 10^{-3}$ | $1.0 \times 10^{-2}$ | N.S. |
| **Olfactory bulb** | | | | N.S. | $9.3 \times 10^{-3}$ |
| **Cerebellum** | | | | | N.S. |
| **Subcortex** | | | | | |
